# A Novel Approach for Using Thermal Imaging to Study Behavioural Responses to Environmental Stressors in Passerines

**DOI:** 10.1101/2024.05.21.595154

**Authors:** Joseph Roy, Ross Macleod

**Affiliations:** School of Biological and Environmental Sciences, Liverpool John Moores University, UK; Scottish Center for Ecology and The Natural Environment, University of Glasgow, UK

## Abstract

Infrared thermography is a promising tool for assessing proxy for stress response in animals. With the advancement of longer recording capabilities, this technique allows the monitoring of the dynamic nature of the temperature variation which is highly correlated to the stress faced by the animal. The current method of extraction of data from thermal videos involves manual identification of the beginning and end of the activity (feeding bout in our case) and extracting the temperature data into CSV files. In this paper, we aim to identify the beginning and ending of the feeding bout by the Identification of Temperature Change (ITC) that happens when a bird arrives and leaves the frame. By identifying this we can monitor longer IRT videos and extract the thermal imaging data for stress monitoring in a faster manner. The identification of the beginning and end frames, also allows us to get behavioural data such as feeding bout length and feeding frequency which also act as a proxy for perceived stress. This method was developed to simplify the data analysis process of thermal imaging videos and proved to be an effective method with wider applications in behavioural monitoring based on temperature change.

## Introduction

Stress is induced when an organism is exposed to an environmental change and the animal tries to get back to equilibrium. Stress can induce several changes in the animal which include behaviour, physiology and demography the animals (Gill, 2007). At the physiological level, under a stressed condition, the circulating glucose, fatty acids, and amino acids are raised. Also, the heart rate and rate of respiration increase. The non-necessary services of the body are suspended and the blood is drawn to the core organs for an emergency response (Jerem et al., 2015). Stress can be acute; which is a shorter high-intensity response or chronic which is a longer-lasting response to a stressful environment. Both kinds of stress can have a detrimental impact on the animal in the longer term and it is important to understand the stressors and the effect of them on animals to alleviate and create a better environment for the animals (refs).

The traditional methods of stress monitoring mostly relied on circulating glucocorticoids which can be considered as a proxy for stress. As they represent the transferred amount of energy in blood, they can also vary with other factors such as age and reproductive status (Jerem et al., 2015). Also, these methods are invasive creating a hurdle in doing the measurement without inducing further stress. For example, in birds, the extraction of blood for stress monitoring involves mist-netting the animals, which are then handled and blood drawn. Often these activities create further stress which can impact the animal negatively as well as create unwanted biases in the results. For methods such as the measurement of heart rate or motion using sensors, often animals have to be attached to animals, which involves handling and adding additional weight to the animal.

There have been significant advancements in recent years in less invasive methods of stress monitoring. Several methods such as blood volume pulse, pupil diameter, salivary glucocorticoid levels, skin conductivity and skin temperature can be used to measure the stress response has been tried successfully (Peña-Bautista et al., 2019; Zhai, 2006). Although, such methods are mostly used in humans and are harder to implement in animals, especially in the wild. In animal wellbeing studies there have been recent advancements using hair and feathers to identify hormonal markers that have shown to be successful in understanding chronic stress (Macbeth et al., 2010; Stubsjøen et al., 2015). Feathers are shown to be successful in longer-term monitoring of stress in birds as well (Bortolotti et al., 2008). These are mostly focused on longer-term stress and also need handling of the animals. This is where thermal imaging emerges as a method for measuring stress that can help alleviate the additional stress caused by measurement as well as it will enable the measurement of acute changes in physical activities in response to stress (Jerem et al., 2015). Skin temperature shows concurrence with hormonal and behavioural patterns (Herborn et al., 2015).

Infrared thermography (IRT/Thermal Imaging) measures the emitted infrared radiation which can be used to calculate temperature. As the core temperature changes under stress, several parts of the skin respond to it in different ways, in humans, the nasal temperature drops while the cheeks stay the same (Herborn et al., 2015). It is important to have bare skin to acquire accurate measurement of skin temperature and in birds, eyes act as a major window of heat exchange which helps in the measurement of temperature changes which correspond to the changes in core body temperature (Herborn et al., 2015; Jerem et al., 2015). However, until recently the methods of measurement mostly focused on taking thermal pictures to acquire a measurement, with some exceptions where they have utilized videos which are often recorded with manual intervention and or spliced to extract the data. When using thermal videos for wild animal monitoring, such manual interventions are necessary to identify the time when the animal has come to the frame and to get the reading for that time. This method is often time-consuming and creates an additional bias in the readings.

With the advent of newer thermal imaging cameras that are capable of independent thermal video recording of hours, we propose having longer recordings by setting up the cameras where the animals are expected to come (eg, feeding station) and we propose using **Idenfification of Temperature Change (ITC)** to idenfity the activity patterns and which helped to streamline the extraction process of thermal data from the videos. This data can be used in the traditional way considering it as a proxy for stress (stress monitoring from now on) while also to use it for behavioural monitoring by understanding the feeding bout length and feeding frequency. This paper is focusing on the behavioural aspect which is a novel approach according to the best of our knowledge.

We utilised an urban-rural gradient study system of Passrines with a fixed feeding station set up with a thermal camera at a fixed distance in every site. We have monitored ∼32 sites as part of the study. As videos are recorded as individual thermal frames, these cameras and accompanying software are capable of generating CSV files which include the the frame number and the corresponding temperature values. We have used the FLIR (model) camera to record the videos and FLIR studio (Model) to track the frames and extract the CSV files with the maximum temperature in the frame, as the eyes act as thermal windows, they create the maximum temperature area in the video. In the generated file we used the increase in temperature to identify when the animal arrives in the frame and the drop in temperature for when the animals leave the frame.

In summary, we assessed each frame in comparison with the previous and the next frame and used identification of temperature change (ITC) to find the points of bird arrival and leaving. The beginning and end frames of each feeding bout which was used to extract thermal data specific to the feeding bouts which was then used for physiological and behavioural analysis.

## Protocol

### Filming Setup and Standardization

1. The filming setup involved a feeding station with a shepherd’s crook with a feeder hanging from it. The feeder was attached using cables to prevent swinging motion. Food was provided ad libitum based based on the manipulation protocols. The feeder also had a black surface with a temperature sensor (tiny tag model) attached to it which was used to calibrate the variation in thermal videos.
2. We kept the thermal imaging camera in an 18L black box which had a hole that pointed to the feeder. The box was also equipped with an acoustic recorder and a monitoring video camera (models) to monitor the level of feed as well as to collect multidimensional data which will be analyzed separately.
3. The opening of the filming box was kept at a 60cm distance from the perch of the feeder which was standardised across sites to account for the distance-based measurement variation in thermal video.
4. The filming was done using a FLIR (model) camera at 15 frames per second. Based on the experimental manipulations for predation risk and food availability, we inserted the thermal camera inside the box and did a one-hour film.

**Figure 1.**
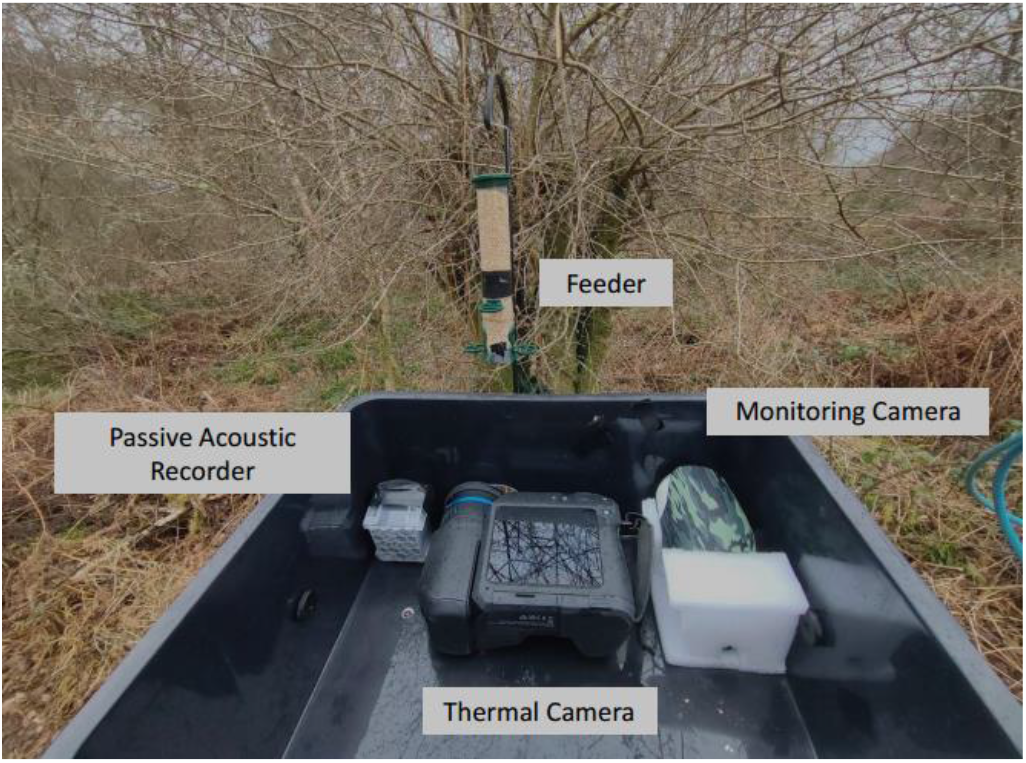
Showing the setup of the multi-input data collection station which we used in the study. There is a combination of a passive acoustic recorder, a thermal imaging camera and a monitoring camera.

### Extraction of Maximum Eye Temperature

1. After filming, the videos were transferred to the FLIR Thermal Studio (model) for processing.
2. We specified the perches as our areas of interest using the rectangle tool and extracted the maximum temperature of the area to represent eye temperature.
3. Also, we extracted the temperature from the standardization black body area for ambient temperature.
4. The data from the thermal video was generated as a CSV file which was then processed in R (R version 4.3.3 (2024-02-29 ucrt, ‘Angel Food Cake’, R Studio Version ‘Ocean Storm’)

**Figure 2.**
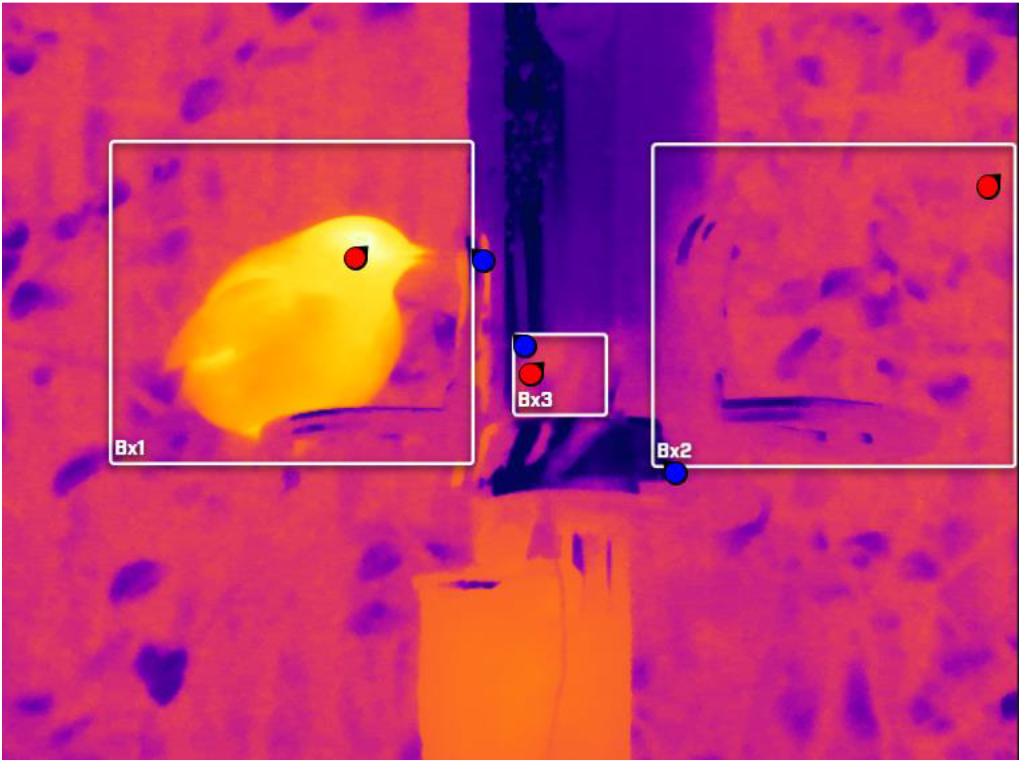
Extraction of Maximum eye temperature from the thermal video using FLIR thermal studio. Box 1 and 2 represent the perch areas that are being measured for the presence of birds. Box 3 is the black body surface which is for the calibration of camera temperature and the measured temperature by the black body.

### Processing the Feeding Bout Data

1. The CSV file extracted from FLIR studio is created for each frame and its corresponding temperature.
2. Identification of temperature change (ITC)-We used a combination of a temperature threshold, along with lag and lead functions in R to identify the changes in temperature in accordance with the previous and next frames.
3. The temperature threshold was implemented using the greater than function and we used 25 degrees Celsius as the threshold to identify the presence of birds. (This can be selected based on the need of the specific case)
4. Using the lag function which subtracts the current row from the previous row and the lead function which identifies the next row to the current row we identified when the bird arrived and left the feeder.
5. Combining the temperature threshold with lag and lead functions we identified the frames which had the bird present in the frame and the data was summarised using the summarise function which to identify the feeding bouts.
6. We only considered bird visits that are more than 3 seconds long for stress analysis, as temperature in the initial stage might be highly connected to flight instead of the perceived stress while on the perch. However, such visits can be used in behavioural level analysis.
7. Although, while using for behavioural analysis such as feeding bout lenth and feeding frequency, it is important to consider all feeding bouts.
8. The filtered data was used for thermal imaging analysis and the feeding bouts for the behaviour analysis.

**Figure 3.**
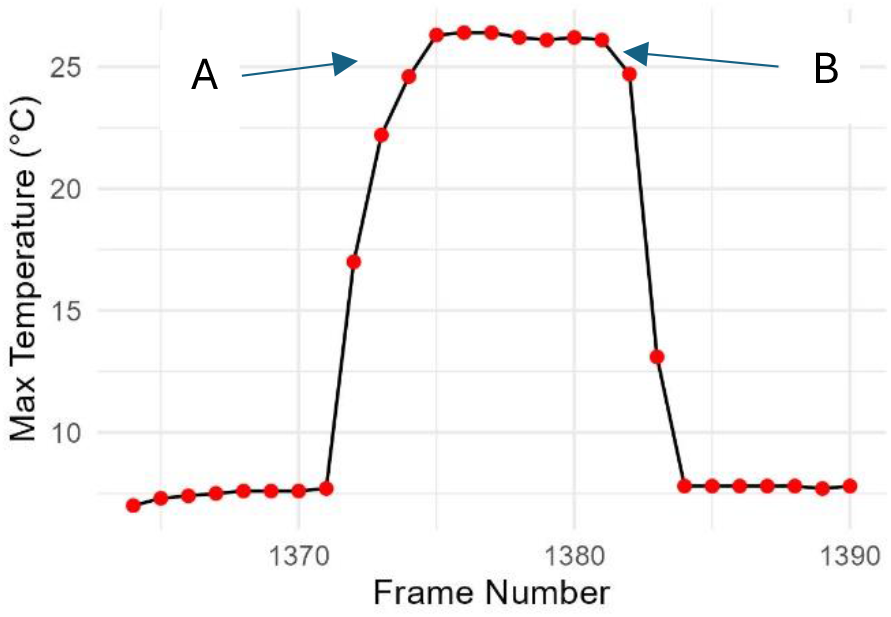
Representation of ITC, this is a figure generated using the CSV file extracted from the FLIR thermal studio software. The graph shows temperature recorded per frame. A-as the bird arrives at the frame, the maximum temperature is raised above the threshold which accounts for the beginning of the feeding bout. B-the drop in temperature denotes the leaving of the bird marking the ending of the feeding bout.

### Representative Results

The major focus of this article is to showcase the identification of temperature change to identify the bird visits which can be used to monitor behaviour characteristics such as feeding bought duration and feeding frequency. Identifying the beginning and end frames using ITC, allows us to identify the frames in which birds are present in the frames which can be used to extract the temperatures to calculate the proxy for stress as done by Jerem et al 2015. As a representation of the methodology, we present results from one urban and one rural site. We are extracting the number of feeder visits and the duration of each visit in one hour, by identifying the temperature change. With the results, we are trying to showcase the method’s effectiveness rather than draw inferences about the study sites.

### Data Extraction

Using the ITC method we identified points of temperature change happening above and below the threshold to identify the bird visit change. Figure 3 represents a graphical representation of temperature change. Using the above mentioned method, we were able to idenify the first and last frames from the CSV files which where then used to generate summary tables as shown in table 1 and 2. The frames corresponding to the visits are extracted from the complete CSV file to be used for the analysis of stress monitoring as mentioned in Jerem et al 2015.

**Figure 3.**
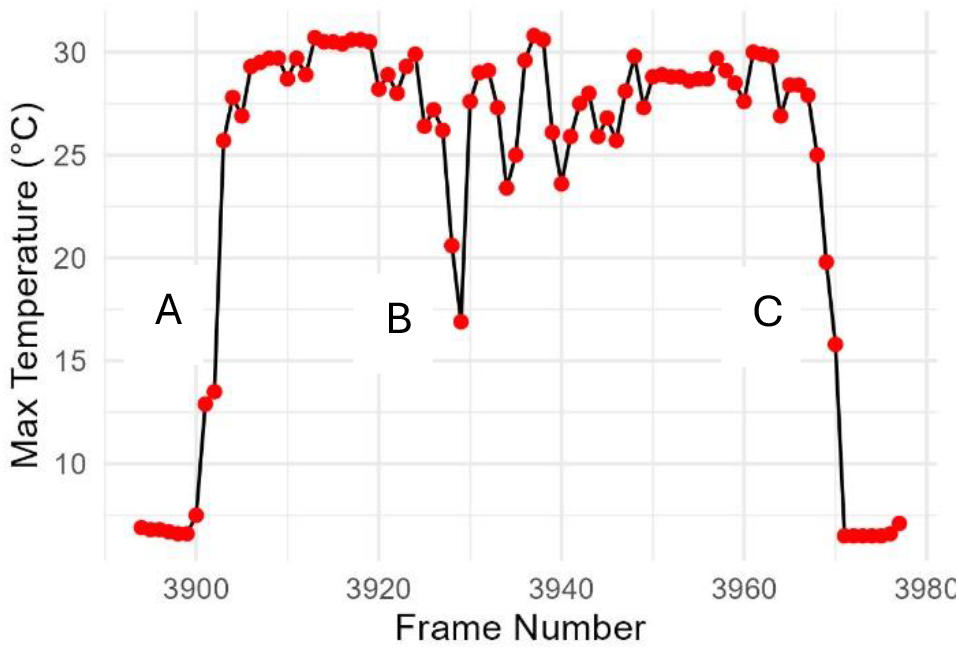
Figure showing the change in maximum temperature when the bird arrives on the feeder and the drop when the bird leaves. Figure shows the feeder visit ID 39 and 34 from table 2, A is when the first feeding bout starts, at B a visit change happens where in a new bird comes and occupy the perch, at C the bird leaves and the temperature drops.

**Table 1 and 2.**
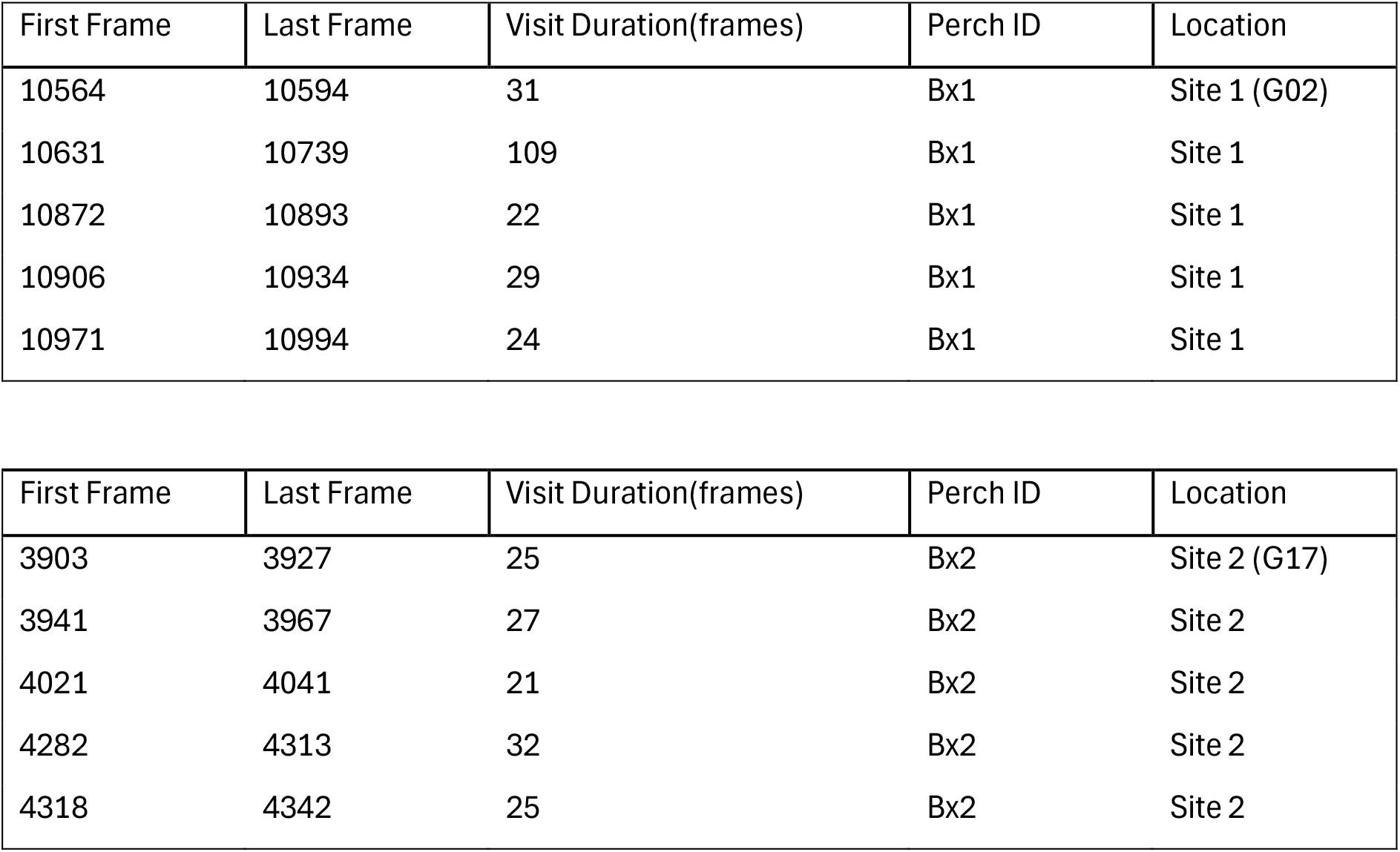
Representation of behaviour data generated by ITC method; By identifying the temperature increase and decrease we can identify beginning and end frames of each feeding bout. This also provides us with the number of frames which can be converted to duration of feeding bout in seconds.

### Generation of Behavioural Data

ITC method is able to generate summary tables as shown in table 1 and 2. This data can be further summarised to generate behavioural data such as feeding bout length and feeding frequency which allows site wise comparisions based on behaviours. Such a comparision is represented in figure 4 and 5.

**Figure 4.**
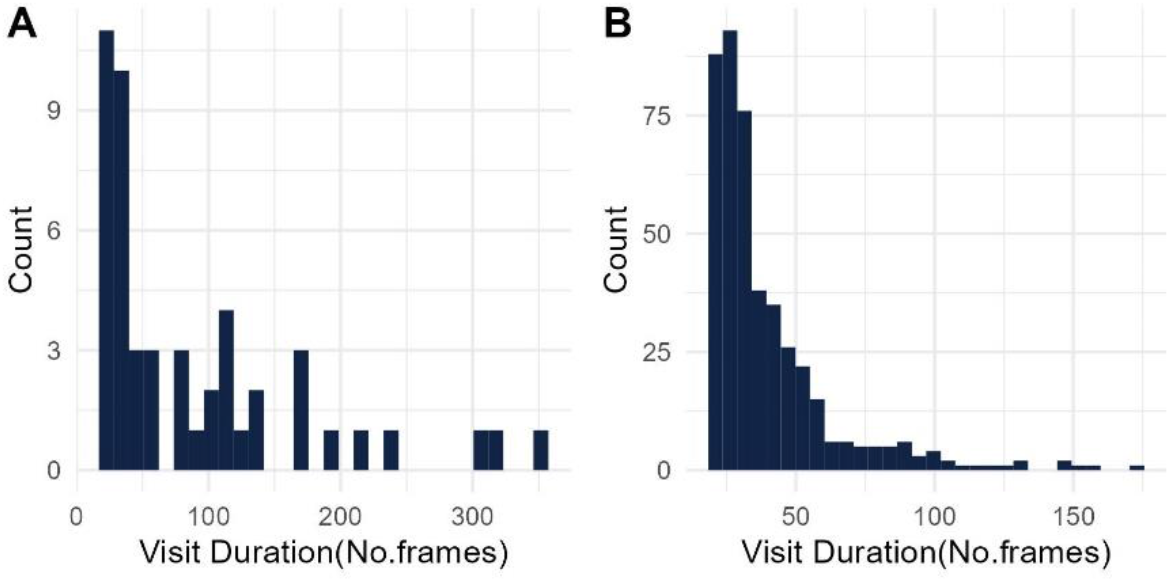
Histograms showing visit duration difference between site 1 and 2. As shown in the graphs, site 1 had fewer visits with longer durations while in site 2 the visits were frequent with shorter length.

**Figure 5.**
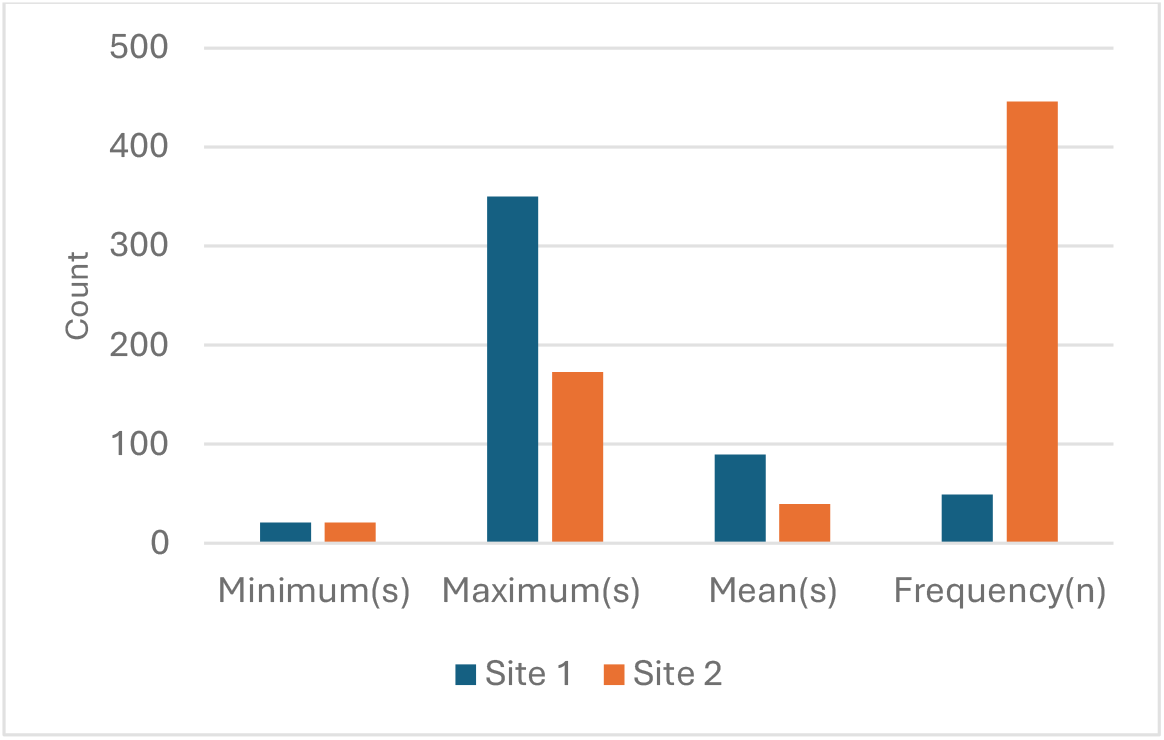
Comparision of behvaioural characters between sites. The graph compares between site 1 and 2. Minimum time of feeder visit is same between two sites, site 1 has the maximum length and the mean of feeding bout is high for site 1 while feeding frequency is remarkably high at site 2.

## Discussion

Thermal imaging is considered as an effective proxy for stress response in animals (Herborn et al., 2015; Jerem et al., 2015). However, the time taken for the extraction of valid thermal imaging data is extremely high which reduces the usability of thermal imaging for longer-term monitoring. Jerem et al., 2015 stated that “the estimated time required to record one bird in the field was about 1 hr, with a further hour required to process the 3 min thermal video sequence”. In our case using the manual method of playing through the videos to identify the beginning and end frame of the feeding bout and extracting corresponding IRT data for the feeding bout data to CSV files took an average of 3 to 6 hours spent per one-hour of recorded video depending on the number of feeding bouts. By using the ITC method we can identify these data in a matter of seconds after the CSV files are provided to R to process. Also, as these are CSV files, they can be processed in batches which allows the processing of large amounts of data in very small amounts of time.

This study aims to simplify the analysis of thermal imaging videos for stress analysis as well as to widen its use for behavioural analysis as well. One of the major aspects of this study is the feeding station-based recording setup which allows standardization of the method across sites. This setup is effective in monitoring stress response in gradient systems in a greater magnitude. Also, the addition of the behavioural data, such as feeding bout length and feeding frequency to the data increases the dimensions in which the stress responses are quantified which allows a clearer picture of the behavioural and physiological aspects of stress response from IRT itself. According to the best of our knowledge, this is the first effort to assess behaviour using thermal imaging. With the combination of a robust setup and the speed of analysis, it proves to be a strong method for IRT-based analysis of the responses of animals to environmental stress.

Species-level identification and demographic characteristics need to be accounted for in analysing this data in our context to get an accurate reading of behavioural data and to avoid species and demographic bias. It is important to select a temperature treshold that is able to accommodate random movements the bird will show inorder to avoid false readings. Also, in our method two perches were present in the frame which creates a need for extraction of the temperature data for each of the perch individually. If the frame only had one perch, the workflow would have been more linear making the extraction process a single event for the whole video. Another aspect to prevent is the presence of another animal in the frame while the perch is occupied. This can be achieved by placing the camera facing perpenticular to the cover/tree. In a controlled environment with only one species, this method can be highly effective in identifying behavioural patterns. It is important to note that the IRT filming needs to be carried out in a standardized manner to get clear and reliable methods. It is extremely important to develop and standardize the experimental setup and to assess the process manually before proceeding to automate the workflow to avoid potential pitfalls.

Aim of this paper is to emphasize the wider applicability of ITC as a strong method of behavioural monitoring in other species as well as other contexts where there is a change in recorded temperature. For example, on a finer level of implementation, the concept of ITC can be used to measure finer aspects of temperature changes. Breathing rate and blinking rate are similar examples. We advise the use of ITC as a concept to develop further the use of thermal imaging for behavioural monitoring rather than relying on the current implementation which is specific for our case. For reference, all the codes are provided in the GitHub repository associated with this paper.

## Notes

### Competing Interest Statement

The authors have declared no competing interest.

